# Snow leopard-mediated apparent competition between wild ungulates and domestic livestock in the Trans-Himalaya

**DOI:** 10.64898/2026.08.05.742978

**Authors:** Deepti Bajaj, Munib Khanyari, Harman Jaggi, Tandup Chhering, Kesang Chunit, Kalzang Gurmet, Tanzin Thinley, Tanzin Thuktan, Rinchen Tobge, Deepshikha Sharma, Charudutt Mishra, Rishi Kumar Sharma, Kulbhushansingh Suryawanshi

**Author notes:** Corresponding author: Deepti Bajaj. Author Contributions: DB, MK, CM, RKS, KS conceptualized the study and CM, RKS, KS, TC, KC, KG, TT, TT, RT formulated the methodology. DB, MK, TC, KC, KG, TT, TT, RT, RKS, KS collected data, while DB, MK, DS, KS curated the data and DB, MK, HJ, KS carried out the formal analysis. MK, TC, KC, KG, TT, TT, RT, DS, KS administered the project, and CM, KS acquired funding and supervised the study. DB wrote the original draft of the manuscript and all authors contributed to reviewing and editing. **Data and materials availability:** Data and code used in this research are available on the Zenodo online repository with the URL: https://doi.org/10.5281/zenodo.21638620 The raw data files for ungulates, snow leopards and livestock have not been uploaded due to safety concerns to avoid misuse (poaching) and can be provided upon request. The data files provided include density estimates of each.

## Abstract

Across High Asia, snow leopards *Panthera uncia* share habitats with pastoralists and consume both wild and domestic herbivores. Increasing wild prey is vital for predator conservation, but its effect on livestock depredation is unclear: it could decrease depredation through apparent facilitation or increase it through apparent competition. We investigated this association using a 13-year time series dataset from Upper Spiti Landscape in the Indian Trans-Himalaya. Long-term data on elusive carnivores are rare, making this time series especially valuable for testing predator-mediated interactions. We found higher wild ungulate density was accompanied by significantly greater depredation of free-ranging livestock, and this association remained positive after bootstrapping to account for uncertainty in density estimates. These results suggest snow leopard-mediated apparent competition between wild and domestic herbivores. We recommend conservation efforts that increase wild prey must also integrate effective livestock protection and support strategies to enhance coexistence between predators, prey and pastoralists.

## 1. Introduction

Across their range, large carnivores often co-occur and interact with humans, with increasing negative interactions due to rising human populations, habitat loss and fragmentation (Cardillo *et al*. 2004; Ripple *et al*. 2014; Woodroffe 2000). A common negative interaction arises from predation on domestic livestock by large carnivores, which results in high economic losses and emotional trauma faced by herders (Dickman 2010; Inskip & Zimmermann 2009), often leading to retaliatory killing of the predator (van Eeden *et al*. 2018; Kissui 2008; Treves & Karanth 2003). Retaliatory killing has led to the extinction of two carnivore species (Woodroffe 2000), and continues to be a major threat to many large terrestrial carnivores worldwide, contributing to their declining populations and contracting ranges (Dickman *et al*. 2011). Facilitating an increase in wild prey populations has been an integral part of large carnivore conservation (Carbone & Gittleman 2002; Karanth *et al*. 2004; Suryawanshi *et al*. 2021b; Wolf & Ripple 2016). However, there is uncertainty on how an increase in wild ungulate density can impact livestock depredation by carnivores.

In a one-predator two-prey system, a higher population of one prey can impact the second prey through two mechanisms. It may be in the form of ‘apparent facilitation’, where an increase in the density of one prey can decrease the predator’s functional response to the second prey (through satiation or switching), which decreases predation of the second prey (Abrams & Matsuda 1996; Long *et al*. 2012). Alternatively, it can be in the form of ‘apparent competition’, where an increase in the density of one prey increases predator density (through a numerical or aggregate response), thereby increasing predation of the second prey (Holt 1977; Long *et al*. 2012; Schmitt 1987).

An increase in wild ungulate density can impact livestock predation by carnivores through similar mechanisms - increases in wild prey density could either decrease livestock depredation through apparent facilitation or increase it through apparent competition. Several studies have shown evidence for apparent facilitation – wolves *Canis lupus* prey less on livestock in areas with higher wild ungulate abundance (Janeiro-Otero *et al*. 2020; Meriggi *et al*. 1996), and several large carnivore species show high livestock depredation where wild prey abundances are low (Khorozyan *et al*. 2015; Soofi *et al*. 2019). However, very few studies have directly investigated and found evidence of apparent competition between wild ungulates and livestock (Suryawanshi *et al*. 2017).

The snow leopard *Panthera uncia*, occurring in the Asian high mountain ranges (McCarthy *et al*. 2017), presents a unique case study to understand these indirect interactions. Snow leopard diet is reported to comprise both wild and domestic prey, with the contribution of livestock ranging from 7% to as high as 65% (Bocci *et al*. 2017; Lu *et al*. 2021; Shehzad *et al*. 2012; Suryawanshi *et al*. 2017). This damage caused by snow leopards to livestock often causes conflict between livestock rearing and snow leopard conservation (Suryawanshi *et al*. 2013) which can drive retaliatory killing, a major threat to its survival (Snow Leopard Network 2014). While population growth of wild prey is recommended for snow leopard conservation (Mishra *et al*. 2003; Shrestha *et al*. 2018; Suryawanshi *et al*. 2021b), its impact on livestock predation has mixed evidence.

Lu et al. (2021) reported the occurrence of apparent facilitation between blue sheep *Pseudois nayaur* and livestock through strong snow leopard diet preference for blue sheep in China. Contrastingly, Suryawanshi et al. (2017), based on data from seven sites, reported apparent competition, and predicted that livestock predation by snow leopards would increase and then stabilize with an increase in wild prey. However, these studies were based on cross-sectional point-data, from multiple study sites monitored over a short period of time. Long-term data allows for a deeper understanding of population dynamics by taking into account temporal heterogeneity and responses that manifest over multiple generations (Karanth *et al*. 2004; Lindenmayer *et al*. 2012, 2022; Reinke *et al*. 2019; Sinclair *et al*. 2007). There is a lack of time series data to understand these dynamics, despite the existence of several long-term monitoring programs for large carnivores (Bakker *et al*. 2020; Harmsen *et al*. 2017; Kelly *et al*. 1998; Nakamura *et al*. 2021; O’Neil *et al*. 2017; Smith *et al*. 2017).

In this study, we investigated the relationship between wild prey density and livestock depredation by snow leopards using a long-term dataset collected over 13 years from the Upper Spiti Landscape (USL) in the Trans-Himalayan region of India. We hypothesized the indirect interactions between wild ungulates and livestock mediated by snow leopards to be apparent facilitation (Fig. 1A), apparent competition (Fig. 1B) or no effect i.e., no indirect interactions between wild ungulates and livestock (Fig. 1C).

**Fig. 1.**
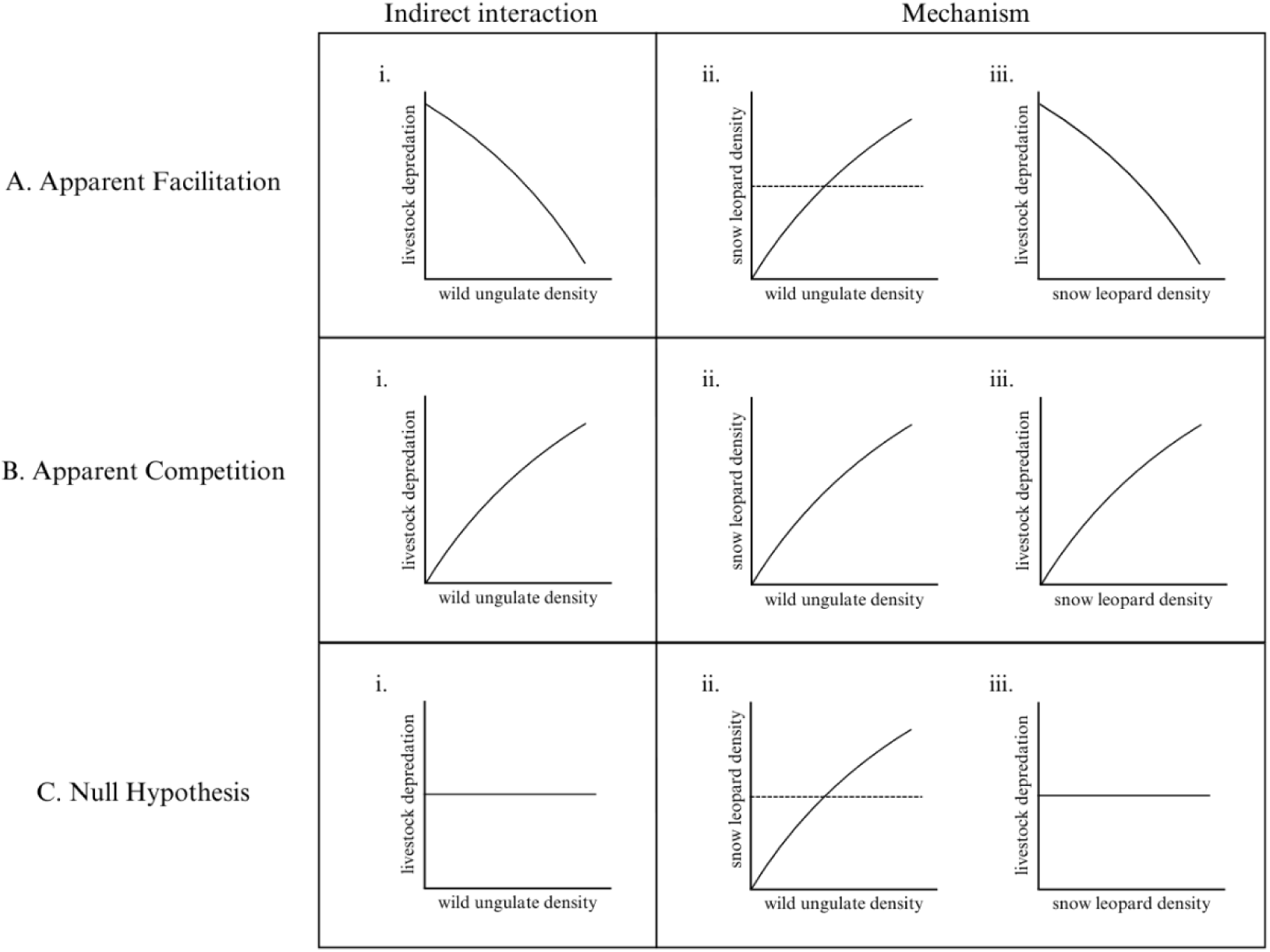
Conceptual diagram of indirect interactions between wild ungulates and livestock mediated by snow leopards in a one-predator two-prey system. These can be in the form of **(A)** apparent facilitation, **(B)** apparent competition or **(C)** null hypothesis, i.e. no indirect interactions between wild ungulates and livestock. In each, **(i)** represents the indirect interaction between wild ungulate density and livestock depredation by snow leopards, while **(ii)** and **(iii)** represent the mechanism of each indirect interaction. This includes **(ii)** the relationship between wild ungulate density and snow leopard density and **(iii)** the association between snow leopard density and livestock depredation by snow leopards.

## 2. Methods

### 2.1. Study Area

We carried out this study in Spiti Valley (c.12,000 km^2^) within the Lahaul-Spiti district of the state of Himachal Pradesh, India. This is a high-altitude region (3500 – 6700 m asl) in the Trans-Himalaya, in the rain shadow of the Greater Himalaya. The vegetation of this region is broadly classified as ‘dry alpine steppe’ (Champion & Seth 1968), and annual temperatures range from −40 °C in peak winter to over 30 °C in summer. For our study, we selected the left bank of the Spiti River, or Upper Spiti Landscape (“USL”; 32.12 - 32.48 °N, 77.73 - 78.27 °E), covering an area of 725 km^2^ (Fig. S1).

The large mammalian fauna of the region comprises wild ungulates such as bharal (blue sheep) and ibex *Capra sibirica*, and their predators, snow leopards, wolves, and free-ranging dogs. The human communities in this landscape are primarily agro-pastoral, practicing seasonal cultivation and livestock rearing. During summer, they grow barley *Hordeum vulgare*, green pea *Pisum sativum*, and its local black pea variety (Jaggi *et al*. 2025). They also rear livestock that graze in the surrounding pastures, except in extreme winters when their diet is supplemented by stall feeding. Livestock here can be categorized into large-bodied free-ranging (“free-ranging”; horse, yak) and medium/small-bodied herded (“herded”; sheep, goat, donkey, cow, cow–yak hybrid).

### 2.2. Data Collection

We collected data on wild ungulates, livestock, and snow leopards annually over 13 years (2011-2023) as part of a long-term monitoring program.

#### 2.2.1. Wild ungulates

We used the double-observer survey method, which is based on mark-recapture theory, to estimate the abundance of wild prey species - blue sheep and ibex (Forsyth & Hickling 1997; Suryawanshi *et al*. 2012). In this method, an ungulate group is the unit that is “marked” and “recaptured” rather than individual animals, since the groups can be identified during the surveys based on various characteristics (Forsyth & Hickling 1997). Two teams surveyed each block on foot, with the second team starting their independent survey 15–30 minutes after the first team (Suryawanshi *et al*. 2012). The surveys involved three main assumptions: (i) entire visual coverage of each block was possible during the survey, (ii) two teams, each with the same number of observers (2-3), conducted the surveys and obtained detections independently of each other and (iii) ungulate groups could be identified individually based on group size, age-sex composition, location and any other peculiarities that the teams could observe. The data collected included the number of groups sighted, group size and detection or non-detection of each group by the two observer teams.

We divided the study area into 21 smaller blocks of ∼30 km^2^ depending on the size and topography of the area, with each block consisting of a set of adjoining sub-catchments, separated from the next block by a ridge. Each survey team independently walked each block within 1.5-3h, annually during late spring (mid-May to mid-June) before the ungulates moved to higher elevations for their birthing pulse, covering a total effort of 189 km each year. The teams, consisting of trained and experienced observers, were constant throughout the years, thereby reducing observer bias.

#### 2.2.2. Livestock

We conducted census via key informant surveys in all 17 villages of USL in December each year (except in 2016 when we were unable to cover five villages due to logistical difficulties), to collect data on livestock population and depredation by snow leopards, wolves, free-ranging dogs, and other causes of death such as diseases like Foot and Mouth Disease (FMD). Key informants included village heads, herders and/or knowledgeable people who were older and more experienced. We surveyed two/three informants from each village each year to triangulate the data, and the same informants were interviewed each year.

Predator kills were identified by key informants based on predator-specific signs (single puncture marks near the neck by snow leopards (Fox *et al*. 2024; Krofel *et al*. 2021), on the flanks/hind by wolves, and multiple marks across the body by packs of free-ranging dogs) and occasionally by direct sightings when the predator would come to feed on the kill. Herder accounts were occasionally verified by field research staff through visits to kill sites. Although we recorded all causes of death, we only used confirmed records of livestock depredation by snow leopards for this study.

Because there is no monetary incentive to attribute livestock depredation to snow leopards, and because this survey is independent and not used to inform the governmental compensation procedure, we do not expect informants to misreport snow leopard kills. Additional details of livestock data collection are provided in supplementary material.

#### 2.2.3. Snow leopards

We collected data on snow leopard density using remotely triggered IR camera traps (Reconyx HyperFire 2/HC500) to systematically sample an area of 725 km^2^. This area is larger than the published home range estimates for snow leopards and is considered adequate to capture multiple home ranges (Johansson *et al*. 2016; Suryawanshi *et al*. 2019). We divided the study area into 5 x 5 km grids and deployed at least one camera trap per grid, with a few additional cameras in some years based on accessibility of locations or availability of functional cameras, resulting in 29-37 camera traps deployed each year (except in 2011 when only 23 camera traps were functional due to logistical difficulties). Each camera station consisted of a single camera mounted at an appropriate height and angle to photograph a passing adult snow leopard’s flank markings. To maximize the probability of detecting snow leopards, we placed the camera traps in the preferred microhabitat within each grid (based on the presence of snow leopard signs) or at sites representing their preferred pathways for movement. We ensured a minimum spacing of 2 km between camera traps to optimize the number of individuals captured while adequately recapturing individuals at different camera traps, as required in SECR designs (Efford & Fewster 2013).

We carried out camera trap-based sampling over a period of 13 years (2011-2023, except in the years 2012, 2014, 2018, and 2019 due to logistical difficulties). Each year, we collected camera trap-based data from late spring into summer (April – September), except in 2011 and 2013 when we sampled in the winter. Cameras operated for a duration of 60 days, a period short enough that an assumption of closure of the population is likely to be met (Alexander *et al*. 2015; Strampelli *et al*. 2022). Details on sampling dates and number of cameras placed each year are provided in Table S4.

### 2.3 Data Analysis

#### 2.3.1. Wild ungulate density

We estimated the abundance of wild ungulates (blue sheep and ibex) in USL using the *multimark* package (McClintock 2015) following Suryawanshi et al. (2021a) in R statistical and programming environment (R Core Team 2024). We estimated wild ungulate density by dividing the estimated abundance by the study area.

#### 2.3.2. Livestock density and depredation by snow leopards

We obtained livestock density for each year by dividing the population of each livestock type by the study area. Because livestock population and density can be influenced by sociocultural factors as well as deaths from other predators and diseases, we used only the number of livestock killed by snow leopards in all further analyses. However, because these factors can affect the availability of livestock to snow leopards, we also repeated all analyses to assess predation risk per livestock by using the livestock population as an offset (Fig S3).

#### 2.3.3. Image processing and snow leopard density

We assigned species tags to all camera trap images using the *digiKam* image management software (https://www.digikam.org/), and we extracted the metadata using the package *camtrapR* (Niedballa *et al*. 2016) in R. Errors in image identification are a major source of bias in population assessments of snow leopards (Johansson *et al*. 2020). Therefore, we used a three-reviewer process for individual identification of snow leopard images to reduce the bias following Suryawanshi et al. (2021b).

We estimated snow leopard density for the different years using a maximum-likelihood based Spatially Explicit Capture-Recapture (SECR) model in the package *secr* (Efford 2023) in R. SECR is a form of hierarchical model that explicitly uses the spatial information of detection locations, with a state model representing the distribution of individual home ranges in the region and an observation model representing the detection probability, based on the assumption that the detection probability decreases as the distance between a detector and an individual’s activity center increases (Efford *et al*. 2009). Density represents the intensity of activity centers within the state space and is expressed as a homogeneous Poisson point process. We modelled observations using a ‘count’ detector, which recorded the number of independent captures for each individual at each camera trap. We used a multi-session SECR model (D ∼ session, lambda_0_ ∼ 1, σ ∼ 1) in which years corresponded to sessions (where each session was assumed to have a closed population) as we expected the density to vary over time. We kept the cumulative hazard of detection (lambda_0_) constant because we used the same camera trap locations each year.

We also kept sigma constant since we did not expect the sex ratio, resource availability, habitat, or any other factor that can change sigma to vary over the years, and to control for a single transient individual drastically changing the sigma in any year. We specified a habitat mask with a buffer of 25 km around the traps and a spacing of 1 km between mask points to create a state space of c. 5510 km^2^ around the trap locations. We assumed the expected number of observations of an individual at a trap to vary as a function of the distance between the trap and the activity center of the individual, following a hazard half-normal detection function.

#### 2.3.4. Testing the association between wild prey density and livestock depredation

We tested whether wild ungulates and livestock showed patterns consistent with apparent facilitation, apparent competition, or no indirect interaction mediated by snow leopards (Fig. 1A–C). For each livestock type, we first related livestock depredation to wild ungulate density using negative binomial regression to account for overdispersed count data (Fig. 1i). We then tested two potential mechanisms: the association between wild ungulate and snow leopard density using linear regression (Fig. 1ii), and the association between snow leopard density and livestock depredation using negative binomial regression (Fig. 1iii).

For depredation counts, we modeled *Y_t_*, the number of depredation events in year *t*, as a negative binomial (NB) random variable with mean *μ_t_* and dispersion parameter *θ*:

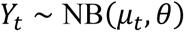

The expected number of depredation events was modeled on the log scale as a function of *X_t_*, where *X_t_* was either wild ungulate or snow leopard density:

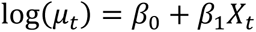

We repeated the above livestock analyses to assess predation risk per livestock by using log(*L_t_*), where *L_t_* is livestock population in year *t*, as an offset term in the negative binomial models:

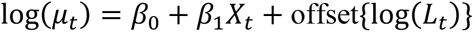

This accounts for the fact that years with more livestock may have more depredation regardless of snow leopard behavior.

To incorporate uncertainty in wild ungulate and snow leopard density estimates, we used a parametric bootstrap for each regression (Efron & Tibshirani 1993; Hutchison *et al*. 2000). For each year, we sampled density values from lognormal distributions parameterized by the estimated mean and standard error, refit the corresponding linear or negative binomial model, and repeated this 10,000 times. We summarized the resulting slope distributions using their means and 95% confidence intervals, treating associations as positive or negative if the 95% confidence interval of the bootstrapped slopes did not overlap zero.

Because predator-mediated interactions may occur with temporal lags, we also tested lagged relationships: wild ungulate density in year (t-1) predicting snow leopard density in year (t), snow leopard density in year (t-1) predicting livestock depredation in year (t), and wild ungulate density in year (t-2) predicting livestock depredation in year (t). The two-year lag allowed for an intermediate response through snow leopard density. Finally, because the time series was short, we conducted a leave-one-out sensitivity analysis on the main regressions without offsets or lags to assess whether results were driven by any single year. Linear models were fitted with lm, and negative binomial models with glm.nb in R.

## 3. Results

### 3.1. Wild ungulate density

The estimated density of wild ungulates (blue sheep and ibex) in USL varied over the study period of 13 years, ranging between 0.94 (95% CI: 0.73 - 1.18) individuals/km^2^ in 2011 to 1.55 (1.24 - 1.89) individuals/km^2^ in 2019 (Fig. 2A, Table S1). The estimated number of groups (Ĝ) varied between 38 in 2011 to 65 in 2021, while the mean group size (µ) ranged between 15 in 2020 to 20 in 2014 and 2015. All estimated parameters for each year are provided in Table S1.

**Fig. 2.**
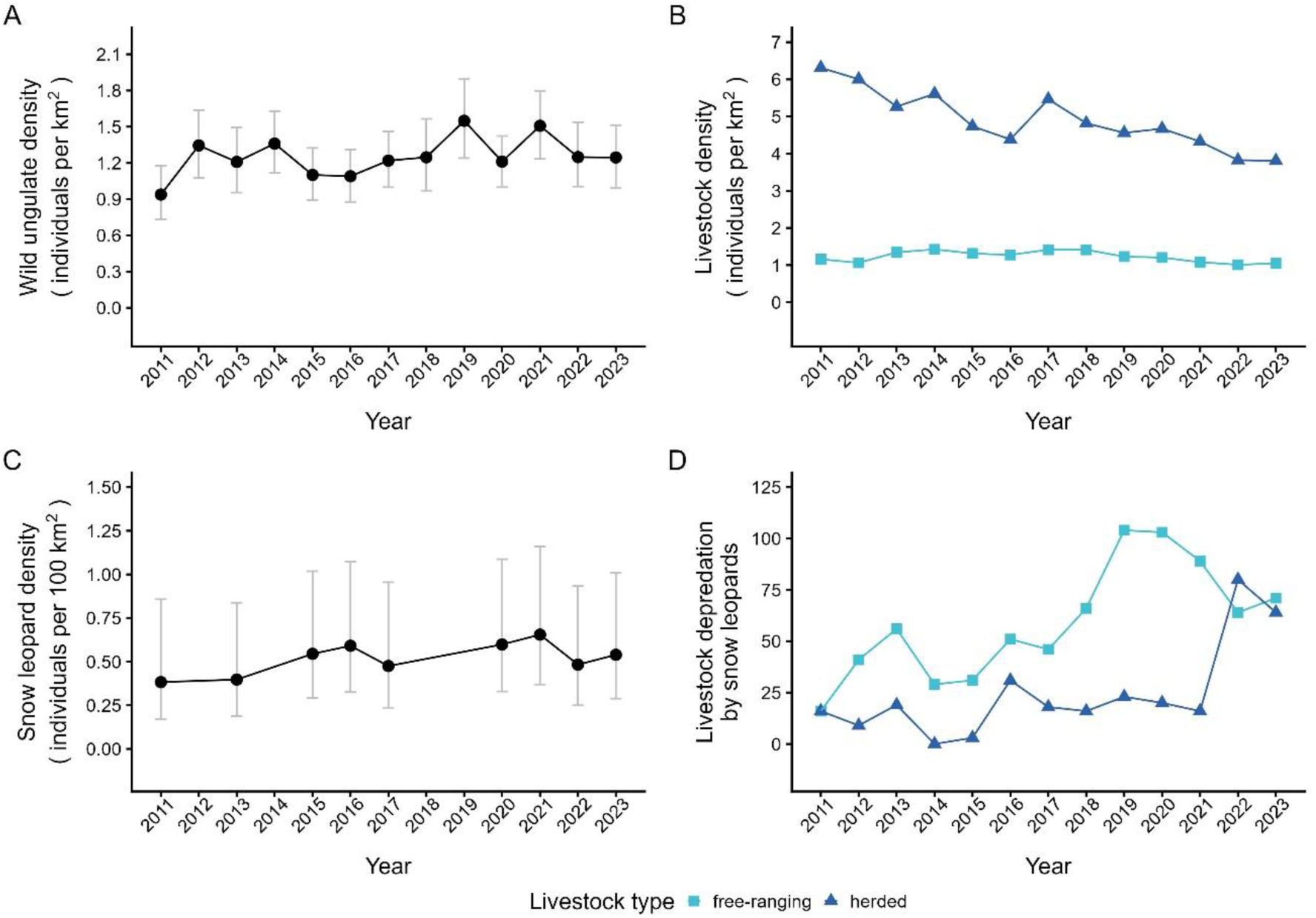
Year-wise trends of density of wild ungulates, snow leopards, and livestock, and livestock depredation for the years 2011-2023 in the Upper Spiti Landscape. This includes (**A**) wild ungulate (blue sheep and ibex) density (individuals/km^2^), (**B**) livestock density (individuals/km^2^), (**C**) snow leopard density (individuals/100 km^2^), and (**D**) livestock depredation by snow leopards. Grey bars in (**A**) and (**C**) represent 95% confidence intervals of the density estimate, colors and shapes in (**B**) and (**D**) indicate the livestock type (free-ranging, herded).

### 3.2. Livestock density and depredation by snow leopards

The density of herded livestock (sheep, goat, donkey, cow, cow–yak hybrid) in all 17 villages in USL decreased over time, ranging between 3.81 individuals/km^2^ in 2023 to 6.31 individuals/km^2^ in 2011. Free-ranging livestock (horse, yak) density was relatively stable, ranging between 1.01 individuals/km^2^ in 2022 to 1.42 individuals/km^2^ in 2014 (Fig. 2B).

Livestock deaths due to wolves, free-ranging dogs, and diseases were higher in herded livestock each year (Table S3), however, snow leopards consistently preyed more on free-ranging livestock (except in 2011 and 2022; Fig. 2D, Table S2). The depredation of herded and free-ranging livestock by snow leopards across all 17 villages in USL showed an increase over time. Depredation by snow leopards of herded livestock ranged from 0 in 2014 to 80 in 2022, and free-ranging livestock ranged between 16 in 2011 to 104 in 2019 (Fig. 2D; Table S2).

### 3.3. Snow leopard density

The overall camera trapping effort comprised 17,169 trap nights in nine annual surveys over a 13-year study period (2011 - 2023) across a state space of c. 5510 km^2^. This resulted in a total of 555 independent snow leopard detections. Snow leopards could be individually identified in 461 (83%) detections. The number of identified snow leopard individuals each year ranged from six in 2011 to 12 in 2021 (Table S4).

Snow leopard density ranged over time between 0.38 (95% CI: 0.17 - 0.86) individuals/100 km^2^ in 2011 to 0.66 (0.37 - 1.16) individuals/100 km^2^ in 2021 (Fig. 2C, Table S3). The cumulative hazard of detection (lambda_0_) was estimated to be 0.028 (0.024 - 0.033), and sigma (σ) was estimated to be 5.0 (4.62 - 5.38) km (Table S4).

### 3.4. Association between wild prey density and livestock depredation

We found that an increase in wild ungulate density was accompanied by a significant increase in the depredation of free-ranging livestock by snow leopards (β = 1.92, SE = 0.67, McFadden’s pseudo-*R^2^* = 0.05, *p* = 0.004). Bootstrap simulations accounting for uncertainty showed a significant mean beta coefficient of 1.18 (95% CI: 0.40 - 2.15), indicating that a 1 individual/km² increase in wild ungulate density was associated with a 3.26-fold (226%) increase in expected free-ranging livestock depredation. However, an increase in wild ungulate density showed no effect on the depredation of herded livestock (β = −0.28, SE = 1.65, McFadden’s pseudo-*R^2^* = 0, *p* = 0.87) with a bootstrapped slope of - 0.13 (−2.15 – 1.83; Figs. 3A, 3B, Table 1). We also found similar results with depredation risk per livestock (Fig. S3), with a significant bootstrapped slope for an increase in wild ungulate density accompanied by an increase in the depredation risk of free-ranging livestock by snow leopards (Figs. S3A, S3B). Our results, thus, suggest snow leopard-mediated apparent competition between wild and domestic prey, specifically free-ranging livestock, with no evidence of apparent facilitation.

**Fig. 3.**
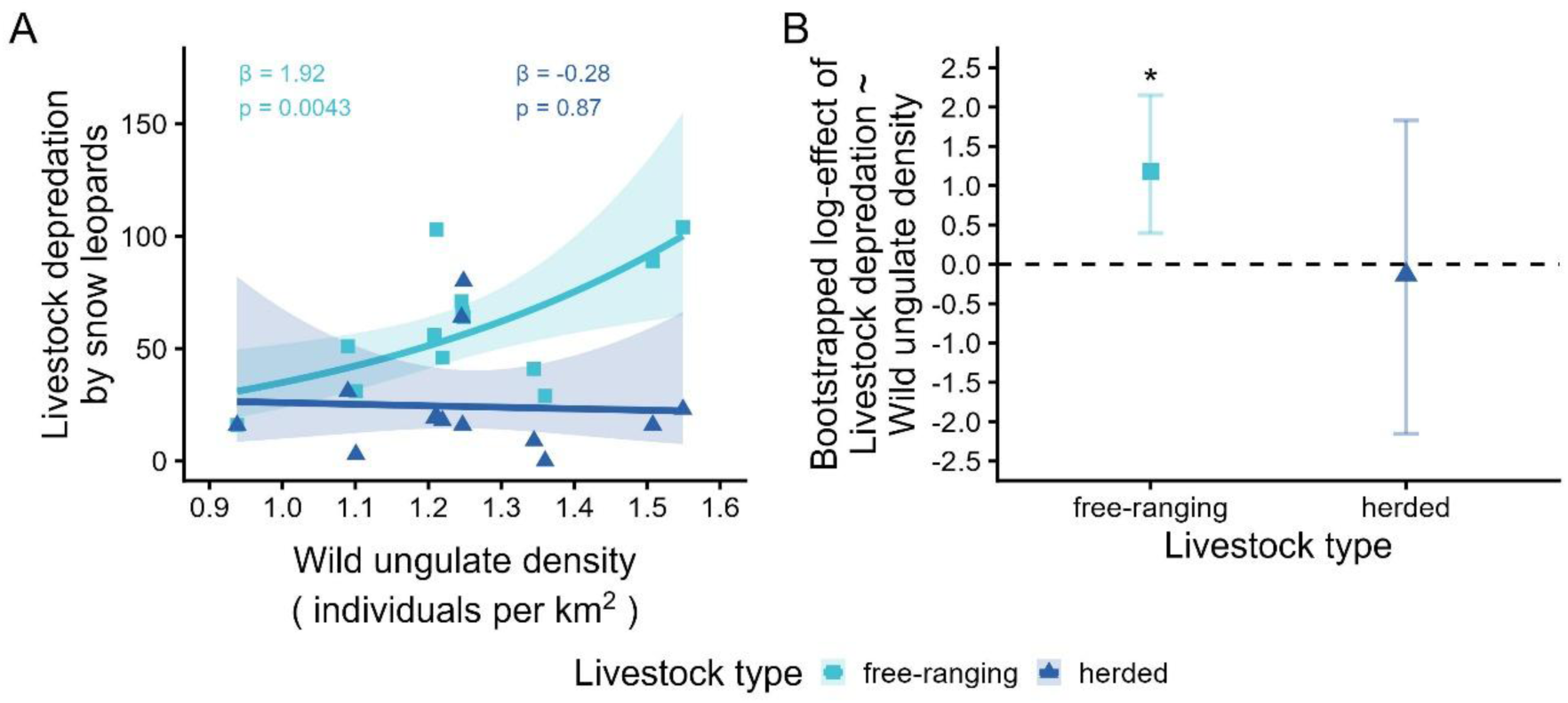
Apparent competition: indirect interactions between wild ungulates and livestock mediated by snow leopards. This includes (**A**) the relationship between wild ungulate density (individuals/km^2^) and livestock depredation by snow leopards fitted with a negative binomial regression, indicating apparent competition. (**B**) shows the mean slope and its 95% confidence interval for the corresponding regression obtained through bootstrapping simulations. Colors and shapes indicate the type of livestock depredated (free-ranging, herded), while shaded regions and error bars indicate 95% confidence intervals. All β and p values have been rounded to two decimal places. Asterix (*) denotes significance.

**Table 1.** Results of regression models and bootstrap simulations examining the relationships between wild ungulate density, snow leopard density, and livestock depredation. β and SE represent the coefficient and standard error from the fitted regression models, with corresponding R² and p-values. Bootstrap mean β and 95% confidence intervals (CI) were derived from bootstrap simulations accounting for uncertainty in density estimates, where relationships were considered statistically significant when the 95% CI did not overlap zero.

| Model | $\beta$ (SE) | $R^2$ | p | Bootstrap mean $\beta$ (95% CI) |
| --- | --- | --- | --- | --- |
| Free-ranging livestock depredation ~ Wild ungulate density | 1.92 (0.67) | 0.05 | 0.004 | 1.18 (0.40 – 2.15) |
| Herded livestock depredation ~ Wild ungulate density | -0.28 (1.65) | 0 | 0.87 | -0.13 (-2.15 – 1.83) |
| Snow leopard density ~ Wild ungulate density* | 0.34 (0.18) | 0.33 | 0.11 | 0.22 (-0.54 – 1.03) |
| Free-ranging livestock depredation ~ Snow leopard density | 3.48 (1.45) | 0.05 | 0.02 | 0.75 (-1.29 – 2.59) |
| Herded livestock depredation ~ Snow leopard density | -0.49 (2.99) | 0 | 0.87 | -1.46 (-3.55 – 3.21) |

\*Linear regression with standard $R^2$ , all other models are negative binomial regressions with McFadden's pseudo- $R^2$ .

To understand the mechanism of this apparent competition, we also found that the relationship between wild ungulate density and snow leopard density was positive (β = 0.34, SE = 0.18, *R^2^* = 0.33, *p* = 0.11; Fig. 4A, Table 1), however, with a non-significant slope of 0.22 (−0.54 – 1.03) through a bootstrapping simulation to account for variation in their density estimates (Fig. 4B, Table 1). Livestock density had a negative effect on snow leopard density (Fig. S2).

**Fig. 4.**
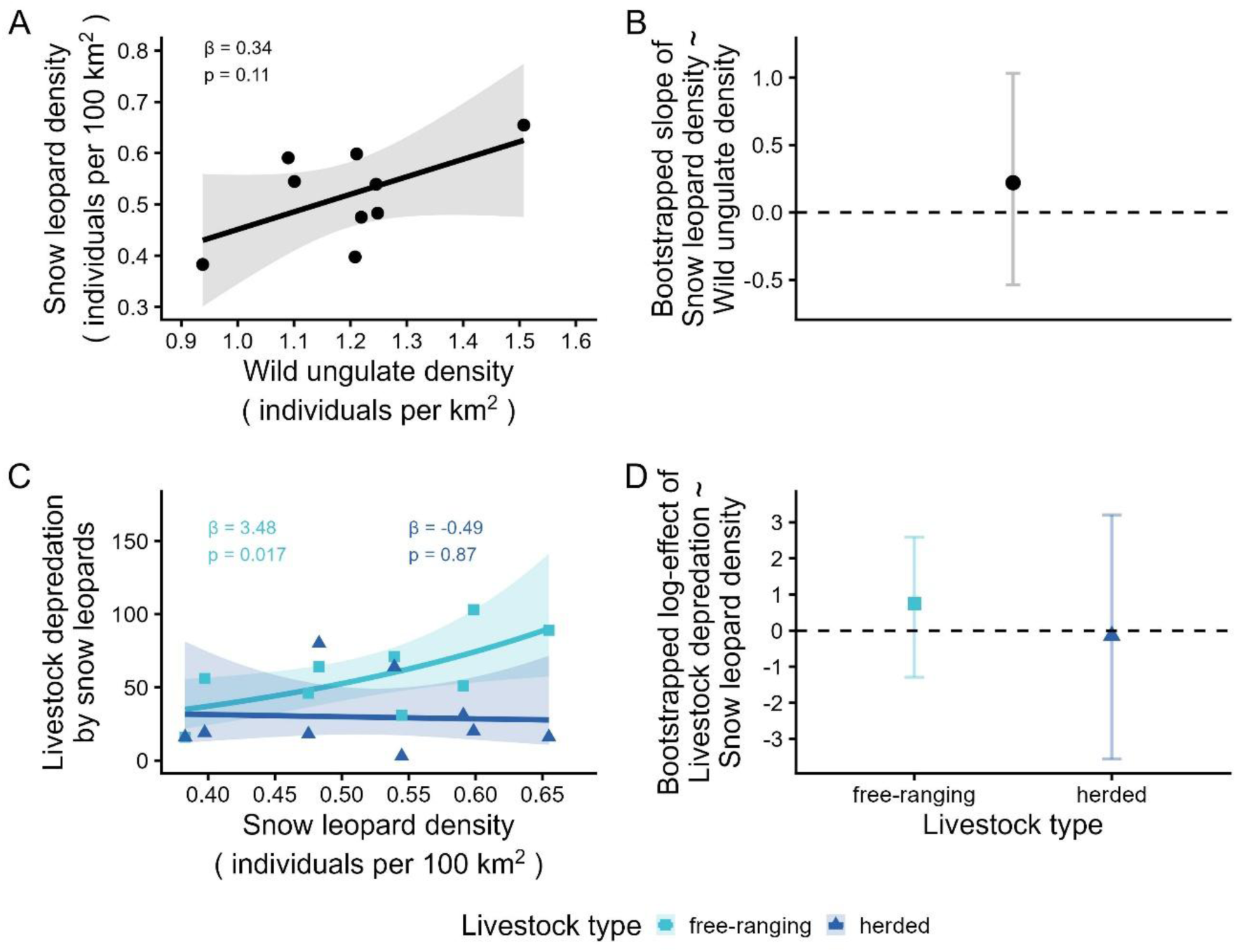
Mechanism underlying apparent competition between wild ungulates and livestock mediated by snow leopards. This includes (**A**) the relationship between wild ungulate density (individuals/km^2^) and snow leopard density (individuals/100 km^2^) fitted with a linear regression, and (**C**) the association between snow leopard density (individuals/100 km^2^) and livestock depredation by snow leopards fitted with a negative binomial regression. (**B**) and (**D**) show the mean slope and its 95% confidence interval for each of the corresponding regressions obtained through bootstrapping simulations. Colors and shapes indicate the type of livestock depredated (free-ranging, herded), while shaded regions and error bars indicate 95% confidence intervals. All β and p values have been rounded to two decimal places.

An increase in snow leopard density was accompanied by an increase in the depredation of free-ranging livestock (β = 3.48, SE = 1.45, McFadden’s pseudo-*R^2^*= 0.05, *p* = 0.02), however with a non-significant bootstrapped slope of 0.75 (−1.29 – 2.59). Increase in snow leopard density had no effect on the depredation of herded livestock (β = −0.49, SE = 2.99, McFadden’s pseudo-*R^2^* = 0, *p* = 0.87) with a bootstrapped slope of −1.46 (−3.55 – 3.21; Figs. 4C, 4D, Table 1), with similar results of depredation risk per livestock (Figs. S3C, S3D).

Indirect interaction models with a temporal lag did not show significance for the effect of wild ungulate density in year (t-2) on livestock depredation by snow leopards in year (t) (Fig. S4A), and for the effect of wild ungulate density in year (t-1) on snow leopard density in year (t) (Fig. S4B). Only the effect of snow leopard density in year (t-1) on free-ranging livestock depredation by snow leopards in year (t) showed a significant increase (Fig. S4C).

The leave-one-out sensitivity analysis carried out on the main results (Figs 3A, 4A, 4C) showed that dropping a year resulted in the beta coefficient of the effect of wild ungulate density on free-ranging livestock depredation varying from 1.27 to 2.14, and on herded livestock depredation varying from −1.00 to 0.21 (Fig S5A). Similarly, the beta coefficient of the effect of wild ungulate density on snow leopard density varied from 0.0057 to 0.42 (Fig S5B). Finally, the beta coefficient of the effect of snow leopard density on free-ranging livestock depredation varied from 2.08 to 5.12, and on herded livestock depredation varied from −2.62 to 1.39 (Fig S5C). In all cases except for those with herded livestock, the direction of the effect remained consistent across dropped years, suggesting the results are not driven by any single influential year.

## 4. Discussion

### 4.1. Snow leopard-mediated apparent competition

Our study suggests apparent competition between wild ungulates and domestic livestock, mediated by snow leopards in the high-altitude landscapes of the Trans-Himalaya. We found that an increase in wild ungulate density was accompanied by a significant increase in free-ranging livestock depredation by snow leopards (Figs. 3A, 3B). While theoretical frameworks of a one-predator-two-prey system predict apparent competition (Abrams & Matsuda 1996; Holt 1977), field evidence from terrestrial carnivore systems has so far been limited. Although our results show statistical non-significance in the mechanism of apparent competition, i.e. in the relationship between wild ungulate density and snow leopard density (Figs. 4A, 4B), and the association between snow leopard density and livestock depredation (Figs. 4C, 4D), the directional signal warrants attention as it may have strong, on-ground implications. These could include increased livestock losses following conservation interventions that increase wild prey, which in turn can lead to negative attitudes towards the predator and retaliatory killing (van Eeden *et al*. 2018; Treves & Karanth 2003). Little is known about indirect interactions between wild predator-prey systems that extend to livestock, despite their management implications for snow leopards and other carnivores that co-occur with human communities.

The positive linear relationship of wild prey density with snow leopard density (Fig. 4A) was consistent with results from other studies on snow leopards and other carnivores (Carbone & Gittleman 2002; Karanth *et al*. 2004; Lu *et al*. 2021; Suryawanshi *et al*. 2017, 2021b). However, the bootstrapping simulations resulted in a weak, nonsignificant positive correlation (Fig. 4B), possibly because of the large confidence intervals in snow leopard density estimates (Fig. 2C). Snow leopards may also show an aggregate response to areas of higher prey densities, where they could achieve higher depredation, rather than a strong numerical response where populations increase linearly with higher prey populations (Cosner *et al*. 1999; Nachman 2006). This is further supported by the non-significance of lagged models (Fig. S4), suggesting an aggregate response within a year rather than a numerical response across years. Snow leopard density also seemed to decrease with livestock density (Fig. S2), although this could reflect shifts in herding practices leading to changes in livestock density (Fig. 2B).

We find a decline in the density of herded livestock (Fig. 2B), largely due to their depredation by free-ranging dogs and wolves, diseases like FMD (Table S3), and the opportunity cost of daily herding. With recent advances in other income avenues such as agriculture, tourism, and road infrastructure development, herders do not prefer rearing herded livestock (Singh *et al*. 2015). However, herders still maintain a relatively stable population of free-ranging livestock (Fig. 2B, Table S2) despite consistent predation by snow leopards (Fig. 2D, Table S2), because these livestock provide benefits such as tilling farms and dung for fuel and also have cultural value encouraged by religious heads (Singh *et al*. 2015). Thus, while herders prefer to rear free-ranging livestock, our data suggest that these livestock face the consequences of apparent competition from wild prey.

We recorded an increase in wild ungulate density accompanied by a significant increase in free-ranging livestock depredation, even after bootstrapping to account for variability in density estimates (Fig. 3A, 3B). These results were consistent when modelling depredation risk per livestock (Fig. S3), suggesting that the association is not driven by variation in livestock availability across years. Although the increase in depredation from 2011 to 2019–2020 seems high (Table S2), it equates to each snow leopard killing three free-ranging livestock in 2011 and nine in 2020 (Table S4). Given that a snow leopard requires about 45 ungulates in a year (Johansson *et al*. 2015), and that livestock may comprise up to 50–60% of its diet (Bocci *et al*. 2017; Suryawanshi *et al*. 2017), this increase is plausible. Therefore, an increase in wild ungulate density could be accompanied by increased free-ranging livestock depredation through a few more snow leopards in the landscape. However, we recorded no effect on herded livestock depredation (Fig. 3A, 3B), likely because herded livestock are grazed as one unit, accompanied by herders, and penned in corrals at the end of the day, whereas free-ranging livestock are left unaccompanied in pastures for several months. We thus suggest that free-ranging livestock may face apparent competition by wild ungulates mediated by snow leopards, and that such interactions may be masked if livestock types with different herding practices are studied together.

Our finding of apparent competition between wild prey and livestock mediated by snow leopards is consistent with Suryawanshi et al. (2013, 2017) but contradictory to Lu et al. (2021), who reported evidence for apparent facilitation. Our study site is a relatively drier landscape than the Sanjiangyuan region in China (annual mean precipitation: 120–500 mm vs. 262–773 mm; (Lu *et al*. 2021)), and possibly driven by bottom-up processes (Mishra 2001). The lower primary productivity (mean aboveground biomass: 170 kg/ha vs. 841– 2880 kg/ha; (Shu *et al*. 2022)) and resultant resource competition with livestock have been reported to limit wild ungulate populations (Mishra 2001; Mishra *et al*. 2004) and dampen blue sheep population cyclicity (Sharma *et al*. 2024). Since wild ungulate availability affects both snow leopard occurrence (Patel *et al*. 2024) and populations (Suryawanshi *et al*. 2017), these bottom-up processes could drive the apparent competition recorded in our study. Moreover, the snow leopard density estimates reported by Lu et al. (2021) rely on encounter rates rather than density estimates derived from spatially explicit capture-recapture, limiting comparison of apparent interaction patterns. Our finding also contradicts Bagchi et al. (2019), who reported decreased livestock predation with wild prey recovery in a partially overlapping study area. However, differences in methodology, including their use of naive counts for prey and predators and lack of genetic confirmation of scats, limit direct comparison. Nevertheless, while our data suggest apparent competition, changes in herding practices, climate change, and increasing human activities may also contribute to variation in livestock and snow leopard trends. These impacts need further investigation.

Few studies have found direct evidence of apparent competition between two prey species mediated by a common carnivore predator. Examples include hartebeest being affected by apparent competition in areas of spatial overlap with zebra mediated by reintroduced lions in Kenya (Ng’weno *et al*. 2019), rodents affecting bird reproduction through shared carnivore predators in temperate forests (Grendelmeier *et al*. 2018), feral horses affecting wild ungulates through shared predators (Boyce & McLoughlin 2021), and higher livestock depredation by lynx in areas of high spatial overlap with natural prey (Odden *et al*. 2008; Stahl *et al*. 2001). Similarly, our study suggests that indirect interactions such as apparent competition can shape predator-prey dynamics involving wild and domestic ungulates.

Finally, we highlight the importance of long-term studies in understanding ecological processes affecting population dynamics, top-down and bottom-up processes shaping populations and assemblages, and guiding evidence-based management (Karanth *et al*. 2004; Lindenmayer *et al*. 2012, 2022; Reinke *et al*. 2019; Sinclair *et al*. 2007).

### 4.2. Management implications

Although an increase in wild ungulate population can support a higher density of snow leopards (Carbone & Gittleman 2002; Karanth *et al*. 2004; Lu *et al*. 2021; Suryawanshi *et al*. 2017, 2021b), the associated increase in free-ranging livestock depredation can increase economic losses for herders and the risk of retaliatory killings. Initiatives supporting higher wild prey populations for the conservation of snow leopards and other carnivores therefore need to be accompanied by context-based conservation interventions and better methods to safeguard livestock and compensate for losses. These can include predator-proof corrals for livestock, especially small-bodied herded livestock, built using local knowledge and incorporating the PARTNERS principles (Bijoor *et al*. 2021; Mishra *et al*. 2017). For free-ranging livestock, measures can include increased vigilance and periodic herding in pastures, as well as better compensation programs including community-led livestock insurance schemes (Mishra *et al*. 2003). Other measures include understanding spatial and seasonal patterns of livestock grazing and depredation sites, and the use of trained guard dogs, deterrents, and barriers to avoid predators (Holland *et al*. 2018). Finally, community-based interventions such as natural resource management programs, education and communication initiatives, local management and monitoring, ecotourism development, and local response teams can raise awareness, build local capacity, and reduce conflict with wildlife (Holland *et al*. 2018).

Our study provides insights into how snow leopard-mediated indirect interactions can affect pastoral communities, as snow leopards are predominantly found outside protected areas in human-use landscapes (Mishra *et al*. 2022). This is particularly important amid a global shift in conservation, away from exclusionary approaches to those that recognize humans and wildlife as parts of an interconnected system (Newing *et al*. 2024). Holistic management practices and continued long-term studies are necessary to build our understanding and maintain coexistence between predators, prey, and pastoralists across mountainous rangeland systems.

## Acknowledgments

This long-term study was carried out in partnership with the Wildlife Wing of the Himachal Pradesh Forest Department and has been possible due to the help of several people over the years. We are thankful to Dorje Angrup, Dorje Bodh, Namgial Bodh, Dorje Chhering, Gonpo Chhering, Tanzin Chhering, Tanzin Dorje, Sonam Lobzang, Dorje Namgial, Tanzin Sherab, Takpa Tanzin, Tanzin Tsewang, and several others who have helped in collecting data over the years. We thank Aakash Lamba, Aditya Malgaonkar, Udayan Rao Pawar, Rajat Rao, Tanaya Rele, Charu Sharma, Aravind Sridharan for snow leopard individual identification, and Ajay Bijoor, Vindhya Jyothi, Abhirup Khara, Abinand Reddy Kodi, Suhel Quader, Devika Rathore, Manvi Sharma, Sidharth Srinivasan and other team members of the Nature Conservation Foundation for insightful discussions and continued support. We also thank the Wildlife Wing of the Himachal Pradesh Forest Department for providing timely permits and continued support. MK would like to thank the European Union - ERC, CONDJUST, 101054259. Views and opinions expressed are however those of the author(s) only and do not necessarily reflect those of the European Union or the European Research Council Executive Agency. Neither the European Union nor the granting authority can be held responsible for them. Lastly, we are also very grateful to the people of Kibber for their warmth and support through the years.

## Funding

Snow Leopard Trust

Wildlife Wing of the Himachal Pradesh Forest Department

National Geographic Society grant ww-067es-17 (KS)

British Ecological Society grant LRB17∖1004 (KS)

