## Supplementary Material for "Snow leopard-mediated apparent competition between wild ungulates and domestic livestock in the Trans-Himalaya"

**This PDF file includes:**

Figs. S1 to S5
Tables S1 to S4

Livestock Data collection:

As livestock is crucial for local livelihoods, villagers periodically check on the well-being of free-ranging livestock. In case of depredation, photographic evidence of the kill marks of the predator involved is obtained, which is crucial to avail compensation. We reported the cause of death as unknown in case herders were unable to confirm the predator identity or if the livestock kill was found after a prolonged duration, and reported as missing if herders were unable to find the livestock. For herded livestock, because people are present with them, eyewitness accounts were recorded.

The government provides equal compensation for livestock killed by snow leopards and wolves, with the compensation amount depending on the market value of the type of livestock and not on the wild predator. However, many herders in the study area do not apply for compensation since the amount received is a small fraction of the livestock value, and the process is time-consuming and often requires travel costs that consume a large proportion of the compensation received.

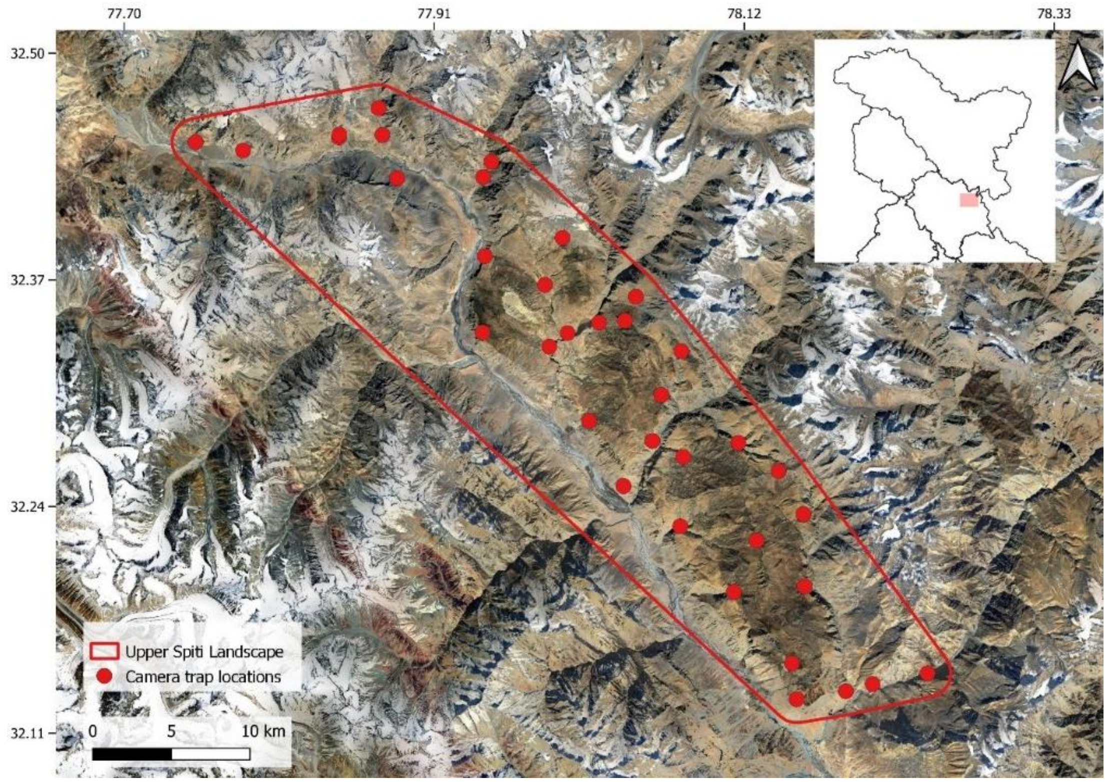


**Figure S1.** Map depicting the Upper Spiti Landscape (USL) and the locations of camera traps each year from 2011-2023.

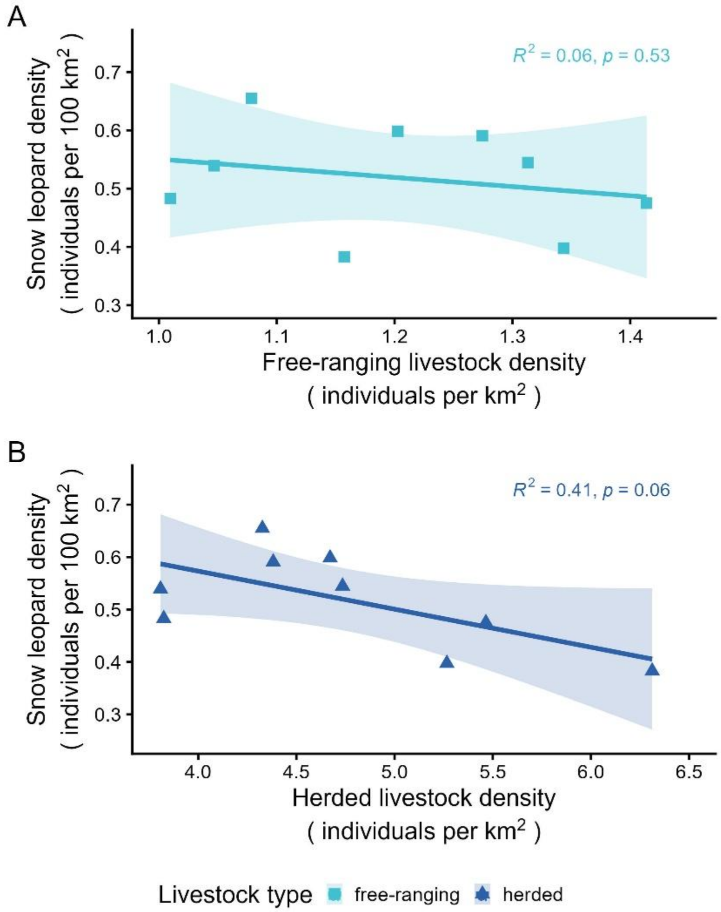


**Figure S2. Relationship between livestock density and snow leopard density**. Here, (**A**) association between free-ranging livestock density (individuals/km$^2$) and snow leopard density (individuals/100 km$^2$), (**B**) association between herded livestock density (individuals/km$^2$) and snow leopard density (individuals/100 km$^2$), both fitted with linear regressions. Colors and shapes indicate the type of livestock (free-ranging, herded), while shaded regions indicate 95% confidence intervals. All $R^2$ and p values have been rounded to two decimal places.

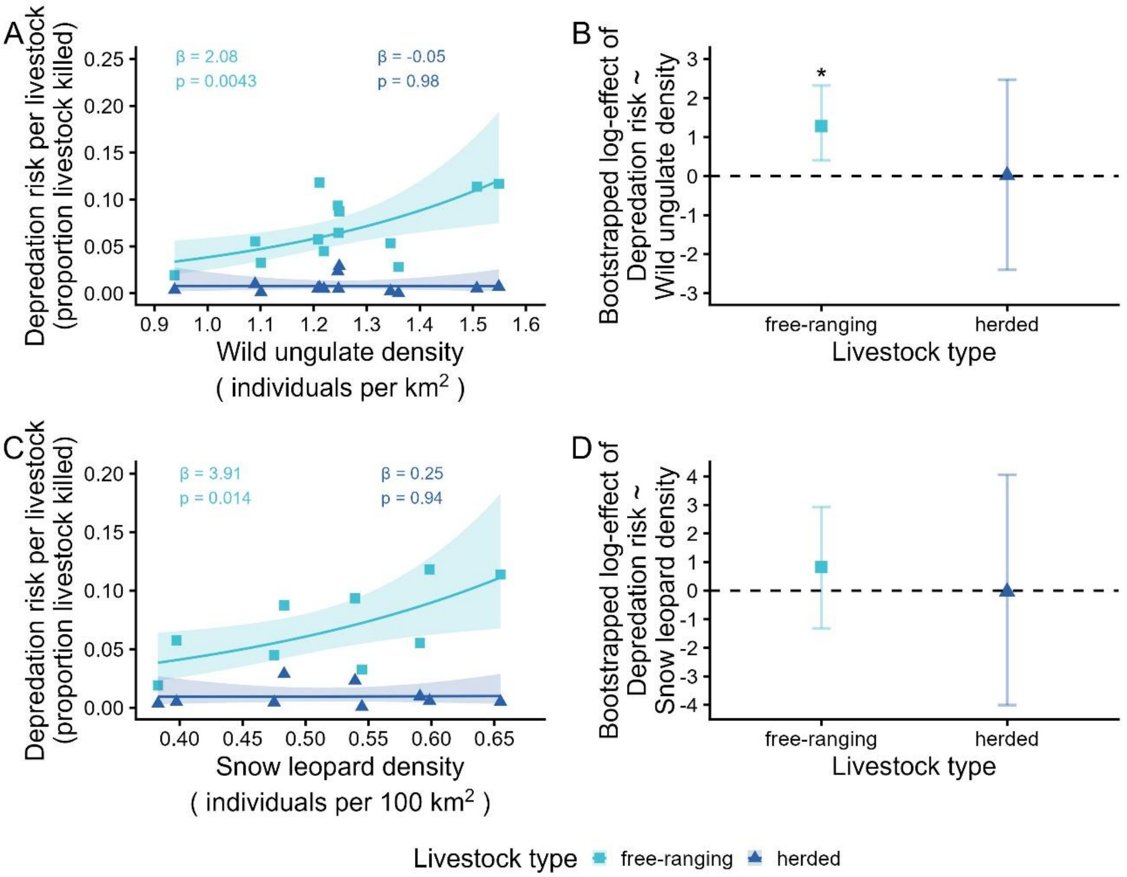


**Figure S3. Apparent competition between wild ungulates and livestock while accounting for livestock availability.** Relationships modelled as depredation risk per livestock by using $\log(L_t)$, where $L_t$ is livestock population in year $t$, as an offset term in the negative binomial model. This includes (**A**) the relationship between increasing wild ungulate density (individuals/km$^2$) and depredation risk per livestock (proportion of livestock killed), indicating apparent competition, and (**C**) the association between snow leopard density (individuals/100 km$^2$) and depredation risk per livestock. (**B, D**) show the mean slope and its 95% confidence interval for each of the corresponding regressions obtained through bootstrapping simulations. Colors and shapes indicate the type of livestock depredated (free-ranging, herded), while shaded regions and error bars indicate 95% confidence intervals. All β and p values have been rounded to two decimal places. Asterix (*) denotes significance.

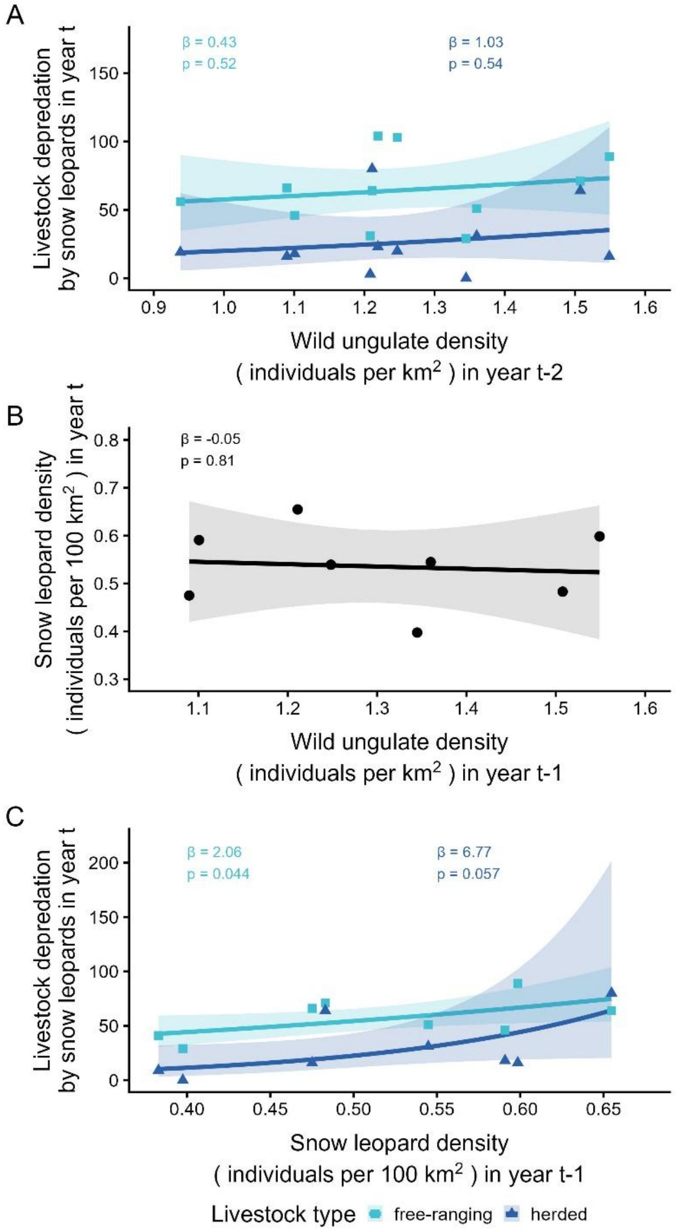


**Figure S4. Assessing indirect interactions with temporal lags.** This includes (**A**) the relationship between wild ungulate density (individuals/km$^2$) in year (t-2) and livestock depredation by snow leopards in year (t) fitted with a negative binomial regression. The mechanism was examined through (**B**) the relationship between wild ungulate density (individuals/km$^2$) in year (t-1) and snow leopard density (individuals/100 km$^2$) in year (t) fitted with a linear regression, and (**C**) the association between snow leopard density (individuals/100 km$^2$) in year (t-1) and livestock depredation by snow leopards in year (t) fitted with a negative binomial regression. Colors and shapes indicate the type of livestock depredated (free-ranging, herded), while shaded regions indicate 95% confidence intervals. All β and p values have been rounded to two decimal places.

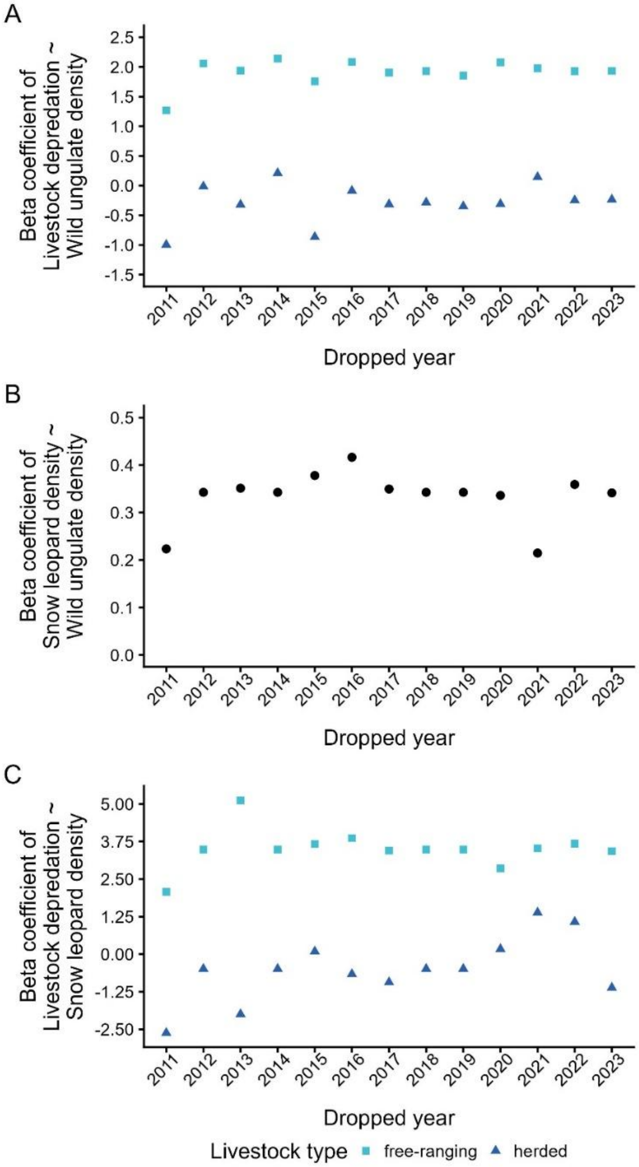


**Figure S5. Leave-one-out sensitivity analysis.** Each point represents the beta coefficient when one year was dropped, assessing the effect of each year on the regression results. Beta coefficients with dropped years for (**A**) the relationship between wild ungulate density (individuals/km$^2$) and livestock depredation by snow leopards fitted with a negative binomial regression (Fig. 3A), (**B**) the relationship between wild ungulate density (individuals/km$^2$) and snow leopard density (individuals/100 km$^2$) fitted with a linear regression (Fig. 4A), and (**C**) the association between snow leopard density (individuals/100 km$^2$) and livestock depredation by snow leopards fitted with a negative binomial regression (Fig. 4C). Colors and shapes indicate the type of livestock depredated (free-ranging, herded).

**Table S1.** Estimated parameters of wild ungulate (blue sheep and ibex) populations using the double observer survey method across the 13-year period (2011–2023) in USL.

| Year | Population size ($N_{est}$) (95% CI) | Population density (95% CI) | Estimated number of groups ($\hat{G}$) | Mean group size ($\mu$) | Detection probability of observer 1 | Detection probability of observer 2 |
|---|---|---|---|---|---|---|
| 2011 | 680 (530 – 853) | 0.94 (0.73 – 1.18) | 38 | 18 | 0.77 | 0.60 |
| 2012 | 975 (781 – 1188) | 1.35 (1.08 – 1.64) | 59 | 17 | 0.74 | 0.70 |
| 2013 | 876 (692 – 1083) | 1.21 (0.96 – 1.50) | 52 | 17 | 0.71 | 0.77 |
| 2014 | 986 (809 – 1179) | 1.36 (1.12 – 1.63) | 48 | 20 | 0.74 | 0.73 |
| 2015 | 798 (646 – 960) | 1.10 (0.89 – 1.32) | 40 | 20 | 0.86 | 0.53 |
| 2016 | 790 (636 – 951) | 1.09 (0.88 – 1.31) | 48 | 16 | 0.87 | 0.55 |
| 2017 | 884 (724 – 1059) | 1.22 (1.00 – 1.46) | 47 | 19 | 0.71 | 0.56 |
| 2018 | 904 (704 – 1135) | 1.25 (0.97 – 1.57) | 51 | 18 | 0.87 | 0.68 |
| 2019 | 1123 (899 – 1373) | 1.55 (1.24 – 1.89) | 61 | 18 | 0.79 | 0.65 |
| 2020 | 878 (724 – 1032) | 1.21 (1.00 – 1.42) | 59 | 15 | 1 | 1 |
| 2021 | 1093 (894 – 1303) | 1.51 (1.23 – 1.80) | 65 | 17 | 0.83 | 0.63 |
| 2022 | 905 (728 – 1115) | 1.25 (1.00 – 1.54) | 51 | 18 | 0.68 | 0.76 |
| 2023 | 903 (721 – 1097) | 1.25 (1.00 – 1.51) | 58 | 16 | 0.86 | 0.53 |

**Table S2.** Livestock population, density (individuals/km$^2$), and livestock depredation and predation rate by snow leopards for the years 2011-2023 collected through census via key informant surveys in all 17 villages of USL in December each year (except in 2016 when we were unable to cover five villages due to logistical difficulties). We calculated predation rate as the number of livestock depredated by snow leopards divided by the population of each livestock type.

| Year | Livestock population | | Livestock density | | Livestock depredation by snow leopards | | Livestock predation rate by snow leopards | |
|---|---|---|---|---|---|---|---|---|
| | Free-ranging | Herded | Free-ranging | Herded | Free-ranging | Herded | Free-ranging | Herded |
| 2011 | 839 | 4576 | 1.16 | 6.31 | 16 | 16 | 0.019 | 0.004 |
| 2012 | 770 | 4352 | 1.06 | 6.00 | 41 | 9 | 0.053 | 0.002 |
| 2013 | 974 | 3818 | 1.34 | 5.27 | 56 | 19 | 0.058 | 0.005 |
| 2014 | 1032 | 4068 | 1.42 | 5.61 | 29 | 0 | 0.028 | 0 |
| 2015 | 952 | 3433 | 1.31 | 4.74 | 31 | 3 | 0.033 | 0.001 |
| 2016 | 924 | 3177 | 1.28 | 4.38 | 51 | 31 | 0.055 | 0.010 |
| 2017 | 1025 | 3962 | 1.41 | 5.47 | 46 | 18 | 0.045 | 0.005 |
| 2018 | 1024 | 3494 | 1.41 | 4.82 | 66 | 16 | 0.065 | 0.005 |
| 2019 | 891 | 3306 | 1.23 | 4.56 | 104 | 23 | 0.117 | 0.007 |
| 2020 | 872 | 3387 | 1.20 | 4.67 | 103 | 20 | 0.118 | 0.006 |
| 2021 | 782 | 3137 | 1.08 | 4.33 | 89 | 16 | 0.114 | 0.005 |
| 2022 | 732 | 2773 | 1.01 | 3.83 | 64 | 80 | 0.087 | 0.029 |
| 2023 | 759 | 2761 | 1.05 | 3.81 | 71 | 64 | 0.094 | 0.023 |

**Table S3.** Data on other livestock deaths, caused due to wolves, free-ranging dogs, disease, and unknown cause or missing livestock for the years 2011-2023 collected through key informant surveys in all 17 villages of USL in December each year (except in 2016 when we were unable to cover five villages due to logistical difficulties).

| Year | Livestock depredation by wolves | | Livestock depredation by free-ranging dogs | | Livestock deaths due to diseases | | Missing livestock/unknown cause of death | |
|---|---|---|---|---|---|---|---|---|
| | Free-ranging | Herded | Free-ranging | Herded | Free-ranging | Herded | Free-ranging | Herded |
| 2011 | 6 | 52 | 0 | 137 | 3 | 28 | 0 | 58 |
| 2012 | 9 | 32 | 0 | 84 | 52 | 28 | 4 | 35 |
| 2013 | 2 | 14 | 0 | 124 | 2 | 21 | 5 | 12 |
| 2014 | 0 | 33 | 19 | 64 | 0 | 14 | 3 | 1 |
| 2015 | 5 | 32 | 25 | 102 | 6 | 66 | 2 | 16 |
| 2016 | 0 | 48 | 14 | 88 | 1 | 15 | 5 | 9 |
| 2017 | 0 | 38 | 4 | 50 | 0 | 32 | 0 | 3 |
| 2018 | 15 | 27 | 0 | 94 | 3 | 38 | 1 | 5 |
| 2019 | 8 | 10 | 14 | 107 | 7 | 47 | 4 | 15 |
| 2020 | 0 | 5 | 2 | 94 | 2 | 24 | 2 | 15 |
| 2021 | 0 | 0 | 2 | 56 | 9 | 274 | 1 | 0 |
| 2022 | 3 | 21 | 8 | 163 | 1 | 32 | 2 | 7 |
| 2023 | 0 | 11 | 5 | 112 | 1 | 26 | 0 | 0 |

**Table S4.** Summary of the camera trapping sampling effort, snow leopard capture details, and estimates of snow leopard density (D; individuals/100 km$^2$) from the multi-session SECR model (D ~ session, $lambda_0$ ~ 1, $\sigma$ ~ 1) across the 13-year period (2011–2023, except in the years 2012, 2014, 2018, and 2019 due to logistical difficulties) in USL. The estimate of cumulative hazard of detection ($lambda_0$) for the same is 0.028 (95% CI: 0.024 - 0.033) and sigma ($\sigma$) is 4984.78 (4619.59 – 5378.83) m.

| Year/ Session | Sampling period (60 occasions each) | No. of camera traps placed | Trap nights | State space (km$^2$) | No. of independent detections identified | No. of independent detections discarded | No. of individuals identified | No. of individuals captured only once | No. of individuals captured at only one location | Density (95% CI) |
|---|---|---|---|---|---|---|---|---|---|---|
| 2011 | 22 Oct - 20 Dec | 23 | 1352 | 5720 | 43 | 13 | 6 | 2 | 2 | 0.38 (0.17 - 0.86) |
| 2013 | 1 Dec 2013 - 29 Jan 2014 | 31 | 1645 | 5643 | 28 | 8 | 7 | 3 | 4 | 0.40 (0.19 - 0.84) |
| 2015 | 1 June - 30 July | 29 | 1709 | 5608 | 43 | 8 | 10 | 1 | 2 | 0.55 (0.29 - 1.02) |
| 2016 | 1 July - 29 Aug | 36 | 2160 | 5615 | 52 | 8 | 11 | 1 | 3 | 0.59 (0.33 - 1.07) |
| 2017 | 3 June - 1 Aug | 30 | 1777 | 5202 | 55 | 16 | 8 | 2 | 3 | 0.48 (0.24 - 0.96) |
| 2020 | 23 July - 20 Sept | 36 | 2091 | 5458 | 71 | 5 | 11 | 0 | 0 | 0.60 (0.33 - 1.09) |
| 2021 | 5 Aug - 3 Oct | 35 | 2065 | 5449 | 66 | 13 | 12 | 1 | 5 | 0.66 (0.37 - 1.16) |
| 2022 | 7 June - 5 Aug | 36 | 2160 | 5449 | 45 | 16 | 9 | 0 | 0 | 0.48 (0.25 - 0.93) |
| 2023 | 7 June - 5 Aug | 37 | 2210 | 5449 | 58 | 7 | 10 | 1 | 2 | 0.54 (0.29 - 1.01) |
